# Insights from the lack of an enigmatic trait: monomorphic sperm in evergreen bagworm moths and the evolution of sperm dimorphism

**DOI:** 10.64898/2025.12.05.692592

**Authors:** Andrew J. Mongue, Thomas W. Johnson, Tracy Liesenfelt, Amanda Markee, Petr Nguyen, Monika Hospodářská

**Affiliations:** University of Florida, Institute of Food and Agricultural Sciences, Department of Entomology and Nematology, Gainesville, FL, USA; Indiana University, Department of Biology, Bloomington, IN, USA; American Museum of Natural History, New York, NY, USA; Biology Centre of the Czech Academy of Sciences, Institute of Entomology, České Budějovice, Czech Republic; University of South Bohemia, Faculty of Science, České Budějovice, Czech Republic

## Abstract

Reproductive traits contain some of the most bizarre and unintuitive innovations seen across the tree of life. Chief among these is the sperm dimorphism of butterflies and moths (Lepidoptera). Males make two types of sperm: traditional fertilizing sperm (eupyrene) and a second, non-fertilizing type (apyrene) that lacks a nucleus entirely. Despite decades of study, the function and evolution of this second sperm type has remained unclear. Here we explore the sperm biology of the evergreen bagworm moth, *Thyridopteryx ephemeraeformis* (Lepidoptera: Tineoidea: Psychidae), and find no evidence for non-fertilizing sperm in this species. We generate genomic, transcriptomic, and proteomic resources to characterize the apparently monomorphic sperm of this species and compare it to that of other Lepidoptera. The fertilizing sperm of the evergreen bagworm shows key differences from other studied species, especially a lack of proteolytic enzymes used to break down sperm bundles in other species. Combining this and other evidence, we offer new hypotheses for the evolution and function of this enigmatic reproductive trait. Notably, early diverging moths appear to produce far less non-fertilizing sperm in general than later diverging Lepidoptera. We infer that non-fertilizing sperm are not necessary in these taxa, perhaps because less robust packaging of fertilizing sperm enables greater mobility than in later diverging moths.

## Introduction

As evolutionary biologists seek to understand universal principles that govern the diversity of life, model species are indispensable tools. Close study of a few species has revealed broadly applicable insights into inheritance, reproductive biology, sex determination, and countless other topics (Maine et al. 1985; Briggs 2002; Hedges 2002; van Wilgenburg et al. 2006; Kiuchi et al. 2014). Indeed, most non-model system comparisons would be impossible without these foundations. Still, clear results from model species can belie the diverse, complicated innovations of other taxa. To truly understand the evolution of life and its myriad adaptations requires addressing the uncommon and strange biology not found in model species. And in terms of bizarre biology, few of life’s core processes are more variable or stranger than those related to reproduction (Bachtrog et al. 2014; Ross et al. 2022). Many traits related to reproduction are immediately eye-catching such as sexual ornaments or weapons of competition that can strain the physiological limits of the organisms that possess them (Husak and Swallow 2011; O’Brien Devin M. et al. 2019). However, equally strange, if harder to observe, is the vast array of microscopic reproductive traits.

Traits are at least as variable at the level of sperm and eggs as they are at larger scales (Tavares-Bastos et al. 2002; Iossa et al. 2016). For sperm in particular, one of the most poorly understood facets of reproductive biology is the widespread occurrence of sperm heteromorphism, in which males reliably make multiple sperm phenotypes (Leviatan and Friedlander 1979; Carcupino et al. 1999; Swallow and Wilkinson 2002; Sasakawa 2009). The most extreme examples are found in Lepidoptera, the butterflies and moths. Males of most species produce both typical fertilizing sperm, called eupyrene, and a second type, called apyrene, which lacks nuclear DNA and cannot fertilize eggs (Meves 1902); apyrene sperm are truly unique to the order Lepidoptera, not found in their sister order, Trichoptera (Friedländer and Friedlander 1983) or other insects.

Despite not being able to perform the one canonical function of sperm, *i.e*. fertilization of eggs, there is strong evidence that apyrene sperm are not merely failures of spermatogenesis. First, their production occurs in a predictable developmental sequence, with eupyrene sperm made by juveniles and apyrene sperm made by adults (Friedlander 1997). Second, they tend to be produced far in excess of eupyrene sperm, with the ratio of apyrene to eupyrene sperm routinely exceeding 10:1 and sometimes reaching as high as 45:1 (Shepherd and Dickinson 2021). Both types are transferred to females during mating (Solensky and Oberhauser 2009) and, most compellingly, the absence of apyrene sperm inhibits fertilization in a model moth (Takemura et al. 2006; Sakai et al. 2019).

Various adaptive functions have been proposed for apyrene sperm based on these observations, ranging from cells specialized in sperm competition to a bizarre nuptial gift of resources (summarized in Swallow and Wilkinson 2002). The best supported ideas are that apyrene sperm help avoid future sperm competition by delaying female remating (Oberhauser 1988; Wedell and Cook 1999) and/or that they capacitate fertilization in some crucial supporting role (Takemura et al. 2006). More specifically, Sakai et al.’s work demonstrated that absence of apyrene sperm caused eupyrene sperm to fail to migrate to the sperm storage organ (spermatheca) from the site of insemination (the bursa copulatrix), suggesting that the infertility is caused by spatial separation of eggs and fertilizing sperm (2019). But this answer, while a key advancement in understanding apyrene sperm, leads to more proximate and ultimate questions. Broadly, how did such a complicated system evolve and more narrowly, what are the molecular mechanisms of the roles played by apyrene and eupyrene sperm?

To begin to answer these questions, we take a multi-faceted approach. First, we reviewed literature on apyrene sperm from the perspective of modern systematic knowledge. Second, based on reports of few-to-no apyrene sperm in a bagworm moth (family: Psychidae; Shepherd and Dickinson 2021), we generated new data in service of the molecular characterization of sperm in bagworm moths. We have undertaken a detailed characterization of sperm in the evergreen bagworm, *Thyridopteryx ephemeraeformis* (Lepidoptera: Tineoidea: Psychidae), using cytogenetics and bioinformatics. On the cytogenetic side we counted chromosomes and visualized sperm morphology via microscopy and staining. On the bioinformatic side, we generated (1) a chromosome-level reference genome for *T. ephemeraeformis*, (2) RNA sequencing from numerous tissues to generate a gene annotation and (3) proteomes of sperm and eggs to characterize the molecular composition of this monomorphic sperm. The protein content of dimorphic sperm has already been characterized in two other lepidopteran species: the monarch butterfly, *Danaus plexippus* (Lepidoptera: Papilionoidea: Nymphalidae) (Whittington et al. 2017), and the Carolina sphinx moth, *Manduca sexta* (Lepidoptera: Bombycoidea: Sphingidae) (Whittington et al. 2015). Our new sperm proteome offers an insight into longer-term conservation of sperm proteins as well as presenting potential functional candidates for future study. Putting all these data together, we suggest new, more specific hypotheses for the evolution and function of apyrene sperm at multiple levels.

## Methods

### Literature review of apyrene sperm research

To contextualize findings for the rest of this study, we started by reviewing recent literature on apyrene sperm. Although this is a standard part of any research undertaking, we wish to be explicit about the inferences we have made to unite the findings from apyrene sperm studies, which are sparse and sometimes disconnected. In particular, one recent study reported numbers and proportions of apyrene sperm across previously unstudied groups (Shepherd and Dickinson 2021), but did so in a phylogenetically agnostic way. We used the phylogenetic relationships inferred by Regier et al. (2015) to place these observations in an evolutionary context.

### Sample collection and sequence generation populations

We generated data from two populations of *T. ephemeraeformis* in two distinct rounds. First, we generated RNA sequencing from bagworms collected from the main campus of the University of Kansas in the spring of 2018 (Lawrence, KS, USA). We brought inactive bags, which contain overwintering, fertilized eggs, up to ambient room temperature (24°C) to break diapause, and reared newly hatched larvae on landscaping juniper, *Juniperus sp*., to adulthood. From this lab colony, we dissected tissue and generated the RNA sequencing described below. This population also served as the material for cytogenetic observations of karyotype and sperm morphology (Supplemental Materials).

In a second round of collection, we obtained second instar larvae from landscaping juniper in Ann Arbor, Michigan in July of 2023. We again reared these insects in laboratory conditions [25 ± 5°C, relative humidity of ~75 (Day)-~50 (night), photoperiod of 16:8 (L:D)] using juniper cuttings to feed them until pupation. At time of pupation, male and female bags are visually distinct enough to sex by eye. Nevertheless, we cut open bags and directly confirmed sex of pupae prior to downstream use. We snap froze 600mg of tissue from 5 male pupae using liquid nitrogen, then stored these samples at −80°C until shipping for Hi-C sequencing (Novogene, Sacramento, CA, USA). Additionally, we extracted DNA from 1 adult male for the primary PacBio HiFi assembly (Arizona Genomics Institute, Tucson, AZ, USA) and 1 adult female for Illumina short-read sequencing to identify the Z chromosome (Novogene, Sacramento, CA, USA). Finally, we allowed the remaining males to mature in order to collect proteomic data. From these adults we dissected sperm for proteomic analysis and again visually assessed the absence of apyrene sperm.

### Sperm cytology

Eupyrene sperm remain in bundles of 256 cells until transfer to the female while apyrene sperm readily dissociate into individual cells even in the male reproductive tract (Friedländer et al. 2005). In practice, this makes it easy to distinguish the two types even under light microscopy; larger eupyrene bundles can be seen individually while apyrene sperm form an indistinct haze of smaller cells. While not conclusive in itself, we first observed dissected seminal vesicles and noted the presence of eupyrene bundles but not individual apyrene cells under a dissecting microscope. Moreover, because apyrene sperm lack a nucleus, they can be definitively identified by staining with DAPI; only eupyrene sperm will fluoresce. To verify our initial observations, we treated the entire contents of a ruptured seminal vesicle with DAPI and visualized the sperm content under a fluorescent microscope. *Gene expression*

We used a TRIzol and Phase Lock gel protocol to extract total RNA. We targeted multiple tissues across both sexes with an initial strategy to cover tissue types broadly, rather than deeply, with two biological replicates. The body cavities of adult females are almost entirely filled with mature eggs, however, and the vitellogenin proteins impeded our spin column-based extractions. As such, there was a higher failure rate for female extractions. Ultimately, we generated the following data, each sourced from single individuals (*i.e*., not pooled): adult male and female heads, adult male thorax, adult testes and ovaries, adult male accessory gland, adult female ovipositor, male and female whole pupae, and unsexed larval silk glands.

### DNA extraction for genome assembly

We extracted high molecular weight DNA from a pupal male using an Omniprep kit (G-Biosciences). We largely followed the kit protocol but modified it in a few important ways. In brief, we used a Powermasher II (Funakoshi) to aid in tissue homogenization, extended proteinase K digestion to be an overnight step, and aided DNA precipitation with 2μl mussel glycogen (20mg/ml) and a 1hr rest period at −20°C.

### Sex chromosome identification

We extracted DNA from the anterior third of a pupal female again using the modified Omniprep extraction protocol described above, then sequenced DNA as 150bp paired Illumina reads to a depth of roughly 30x coverage with Novogene’s services to generate data from the heterogametic (Z0) sex. After finalizing our assembly (described below), we aligned these reads to the reference and calculated depth of coverage on each chromosomal scaffold with the coverage command of SAMtools v1.9 (Homer et al. 2009). Because both sexes are diploid for the autosomes, but females are haploid for the Z, we identified the Z chromosome based on its coverage difference between the sexes, a method we have previously successfully deployed multiple times (Mongue et al. 2017; Mongue and Baird 2024).

### Assembly and annotation

We followed a pipeline that has previously given us success in generating chromosome-level assemblies in other insects (Mongue, Markee, et al. 2024; Mongue, Ross, et al. 2024; Ross et al. 2024; Liesenfelt et al. 2025). To begin, we characterized the genome in two ways. First, by means of flow cytometry. We sent snap frozen adult males to the Johnston cytogenetics laboratory at Texas A&M University; the core ran two technical replicates from nuclei obtained from heads to generate a genome size estimate. Second, we generated an *in silico* size estimate by counting k-mers (-m 21) with Jellyfish v2.3.0 (Marcais and Kingsford 2012) to obtain estimates of genome size and sequencing depth from the PacBio HiFi reads. We then assembled these reads with Hifiasm v1.6.1 (Cheng et al. 2021), using the -l 3 parameter for more aggressive haplotig purging and manually setting the expected homozygous coverage based on our k-mer-based calculations (--hom-cov 58). We collected assembly statistics from this first assembly using a custom python script (available on the git repo for this project) and BUSCO v4.1.4 (Manni et al. 2021) using the lepidoptera_odb10 dataset of expected orthologs.

Next we used Arima’s recommended pipeline (described in detail on their git repo: https://github.com/ArimaGenomics/mapping_pipeline/) to align Hi-C reads to the primary assembly. From there, we used YaHS v1.1 (Zhou et al. 2023) to scaffold the assembly automatically. We then visualized Hi-C contacts and made manual modifications using Juicebox v2.17 (Durand et al. 2016). In total, we identified three likely redundant sequences in chromosomal scaffolds that we moved to the debris. We performed a final BUSCO and assembly statistic calculation and moved on to annotation.

We began annotation by masking repetitive sequences in the genome using Earl Grey (Baril et al. 2024), a complete repeat modeling and masking pipeline. We supplied an initial repeat library that combined the general lepidopteran repeat dataset from dfam (Hubley et al. 2016) with annotations from *Eumetea japonica*, one of the few other sequenced bagworm genomes (Yoshioka et al. 2019). Using our masked assembly, we aligned all of our RNA sequencing to the genome using HISAT2 v2.2.1 (Kim et al. 2019), then used these alignments as evidence for BRAKER v2.1.6 (Brůna et al. 2021).

Separately, we also tested a novel *ab initio* approach to annotation. We used Helixer, a machine learning tool, that requires no RNA evidence to make gene predictions (Holst et al. 2023). We supplied the tool with the soft masked assembly and used the *invertebrate* set of initial model weights. We again used BUSCO (Manni et al. 2021) to assess completeness of the annotation. We selected the more complete of the two annotations for downstream analyses.

### Proteomics

We dissected seminal vesicles from adult male *T. ephemeraeformis* and collected sperm from the vesicle with a 20μl pipette. We snap froze collected sperm in molecular grade water until further analysis. We sent these samples to the IDeA National Resource for Quantitative Proteomics (Little Rock, Arkansas, USA); in total we sent 5 biological replicates, each from an individual male. At the core, they performed mass spectrometry with an Orbitrap Eclipse Tribrid and completed a database search against our BRAKER gene annotation using MaxQuant (Max Planck Institute). They set cutoffs for <1% false discovery and 2+ identified peptides to establish protein identity and returned a list of genes found in the sperm. We also generated an egg proteome by dissecting unfertilized eggs from three virgin adult females. We sent eggs to the University of Florida’s Interdisciplinary Center for Biotechnology Research (ICBR) core facility. ICBR also performed mass spectrometry and a database search against the BRAKER annotation using the software PEAKS. Using these two datasets, we classified *T. ephemeraeformis* genes as either belonging to sperm, egg, both proteomes, or neither. We explored predictions of sex chromosome theory by testing for a biased distribution of genes encoding these categories of reproductive proteins with a X^2^ test of independence between the Z and autosomes.

To further investigate the molecular composition of *T. ephemeraeformis* sperm, we compared genes identified in our proteomics datasets to those of the two other Lepidoptera with sequenced sperm proteomes, *Danaus plexippus* (Whittington et al. 2017) and *Manduca sexta* (Whittington et al. 2015). For consistency with published literature and to take full advantage of existing data, we used the same gene annotation versions as in the previous sperm proteomic studies, namely DPOGS2 for *D. plexippus* and Msex2 for *M. sexta* (Whittington et al. 2015; Whittington et al. 2017; Whittington et al. 2019). We drew from lists of proteins found in apyrene or eupyrene sperm (not mutually exclusive) in each species. Separately, we searched for orthologs between all three species using Proteinortho v5.16 (Lechner et al. 2011). From this list we selected 1-to-1-to-1 orthologs as most likely to retain conserved function between species. We then intersected these orthology data with the proteomic categorizations in each species to create summary statistics and identify candidate genes for further study. We tested for differences in the rate of conservation of reproductive proteins across species via a X^2^ test of independence. In terms of candidate genes, we were particularly interested in (1) orthologs identified in the sperm proteomes of all three species, especially those from apyrene sperm in *D. plexippus* and/or *M. sexta* and (2) genes unique to the sperm of *T. ephemeraeformis* or (3) present in *D. plexippus* and M. sexta but missing in *T. ephemeraeformis*.

## Results

### Evolution of apyrene sperm in early Lepidoptera

By placing Shepherd and Dickinson’s observations (2021) of the presence and abundance of apyrene sperm across species into the phylogenetic framework of Regier *et al*. (2015), we recover the evolution of apyrene sperm as very early in Lepidoptera; however, in most early-diverging lineages, where quantitative data are available, eupyrene sperm almost always outnumber apyrene sperm (**Figure 1, left**), except in one taxon with dubious identification.

**Figure 1.**
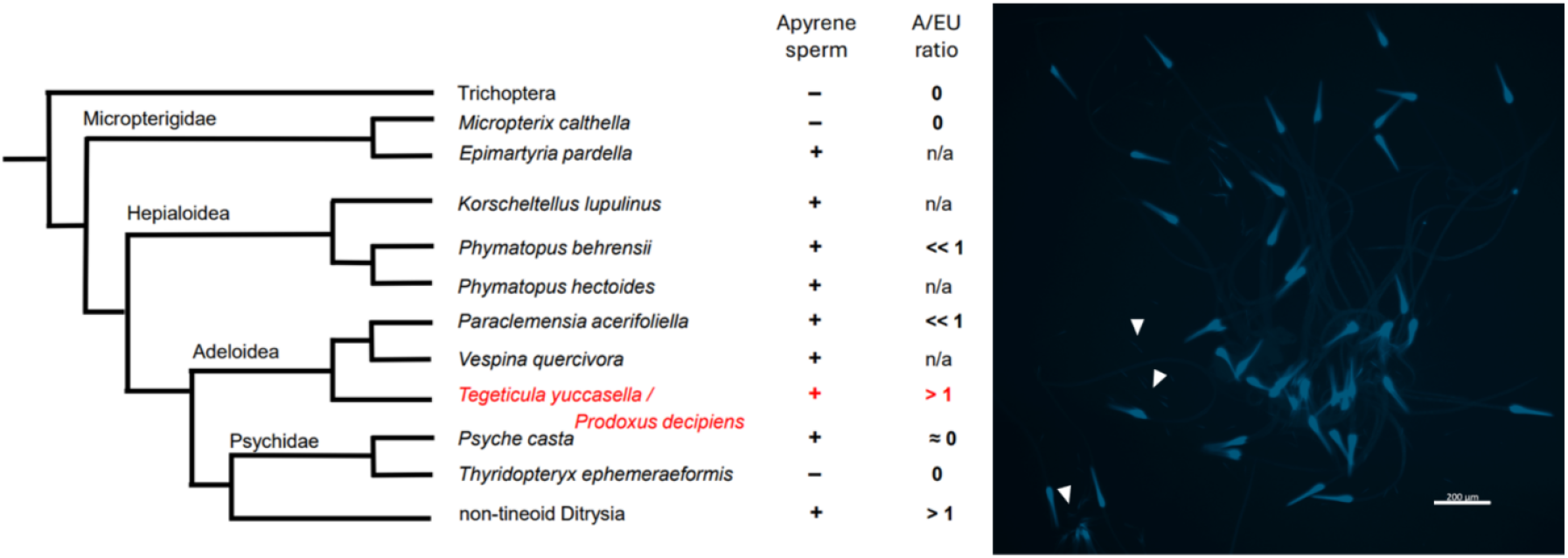
Phylogenetic patterns of apyrene sperm presence and microscopic characterization of sperm in *T. ephemeraeformis*. **Left:** Patterns of apyrene sperm presence and abundance using phylogenetic relationships from Regier et al. (2015) and sperm observations from Shepherd and Dickinson (2021) as well as our own observations in *T. ephemeraeformis*. A/EU ratio is the reported ratio of apyrene to eupyrene sperm, where available (n/a for taxa with no counts). The ambiguity in *Tegeticula/Prodoxus* (in red) represents the original authors’ own uncertainty of the identification. **Right:** Dissected sperm from male *T. ephemeraeformis* seminal ducts shows blue fluorescence when stained with DAPI. The vast majority of sperm are bundled, eupyrene sperm. Even single cells show the presence of a nucleus (smaller light blue rods marked with white arrows), confirming that they are individual eupyrene sperm that have dissociated from a bundle during the staining process.

### Sperm cytology and biology

We collected sperm directly from the adult male reproductive tract, observed its gross morphology, and stained it with DAPI to label nuclear DNA. We found almost exclusively bundles of sperm, as expected of nucleated, eupyrene sperm. Furthermore, these bundles fluoresced when stained with DAPI (**Figure 1, S1 right**), confirming the presence of nuclear DNA. The few single sperm we observed all showed fluorescing nuclei, indicating that they were individual eupyrene cells as well (**Figure 1**). These results strongly suggest a lack of apyrene sperm; although we cannot completely rule out the presence of any apyrene sperm, the fact that none were detected suggests that if they did exist, they were vanishingly rare, in stark contrast to most other Lepidoptera where these data are reported. To further characterize these sperm, we sampled sperm from 5 individual males to create biological replicates for the proteomics dataset described below.

### Sequencing and genome assembly

We generated 24.4 Gb of PacBio HiFi data with a mean read length of 14Kb; accessions for these and all other data can be found in **Table S1**. Subsequent k-mer based analyses indicated that we sequenced to an average depth of 58x homozygous coverage (and 29x for heterozygous regions), and that the genome was roughly 780 Mb. That estimate agrees with male genome size 1C = 790.2 Mb +/-1.6 Mb SE determined by flow cytometry. We assembled a genome slightly larger than these estimates (818 Mb, **Table S2**).

Separately, we generated 17.3 Gb of Hi-C data and 12.6. Gb of Illumina re-sequencing from a female. After manual curation, 98% of the sequence was assigned to the 31 largest scaffolds. These scaffolds, which are substantially larger than the rest in the assembly, correspond to the haploid chromosome count, n = 31, as confirmed by karyotype analysis (**Figure S1**, left).

### Annotation and differential gene expression

We also generated 17 RNAseq datasets, totaling 78.7 Gb of data (**Table S1**). Using these data as evidence for an annotation with BRAKER, we annotated 30,147 protein coding genes, which is almost certainly an overestimate of protein coding genes given results from other Lepidoptera (e.g., ~16,000 in *Danaus plexippus*: Ranz et al. 2021), but it gave high BUSCO completeness with low percentage of duplicates (93.5% complete: 91.2% complete single copy, 2.3% duplicated, 1.2% fragmented, and 5.3% missing). In contrast, the Helixer *ab initio* annotation produced a more reasonable 16,326 protein coding genes, but with substantially lower BUSCO scores (85.6% complete: 83.3% complete single copy, 2.3% duplicated, 4% fragmented, and 10.4% missing). Given the two options, we selected the more complete annotation under the logic that erroneous gene annotations will not show expression in RNA analyses but that the under-annotation clearly excludes valid genes. Indeed, a simple filter for genes with least 5 transcripts per million (summed across all tissue expression) lowered the gene passing gene count to 17,366. Hopefully this comparison proves useful as a test of a modern *ab initio* annotation tool that is still a pre-print (Holst et al. 2023).

### Thyridopteryx ephemeraeformis *sperm and egg proteomes*

Database searching of the full annotation above recovered 2,290 proteins in the sperm; however, presence in sperm does not on its own imply that a protein is sperm-specific. To find sperm-specific proteins, we also generated an egg proteome which contained 1,419 proteins. Intersecting the two, we found 1,070 proteins shared between the two gamete types, leaving 1,220 proteins unique to sperm and a mere 349 unique to eggs (**Figure 2 Top**). A full list of these proteins is available as supplemental data.

**Figure 2.**
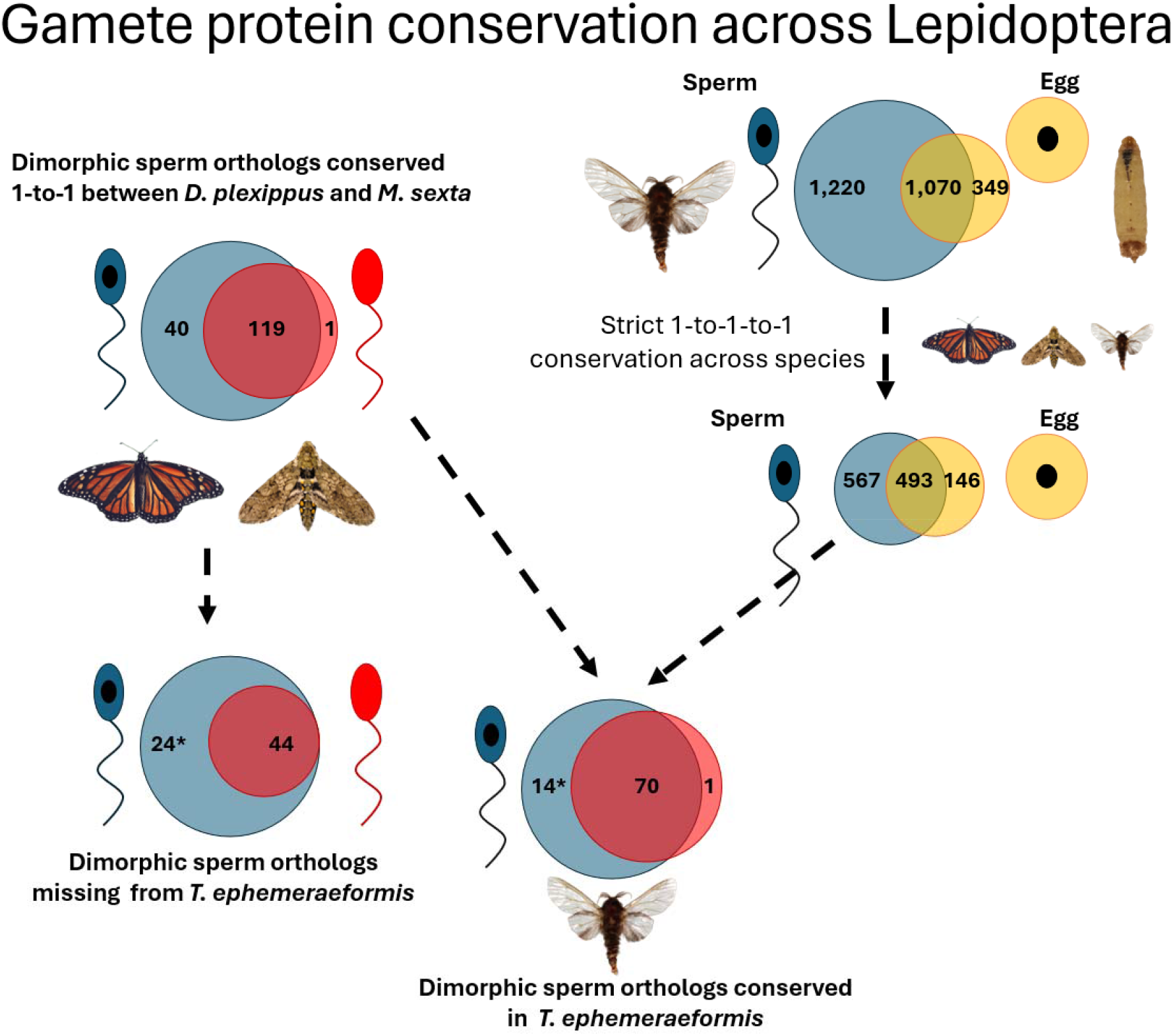
Sperm protein conservation across Lepidoptera with available proteomic data. Where applicable, in each overlap eupyrene proteins are on the left and colored blue, with apyrene on the right in light red and proteins shared between the two morphs in dark red. For sperm and egg comparisons, sperm is on the left in blue with egg proteins on the right in yellow. **Top right:** Composition of the sperm and egg proteomes and their overlap in *T. ephemeraeformis*. **Middle left:** Comparison of 1-to-1 orthologs between *D. plexippus* and *M. sexta*. The most proteins are conserved in the shared portion, followed by eupyrene, with only a single 1-to-1 ortholog found in the apyrene sperm of both species. **Bottom left:** Orthologs found in the two dimorphic sperm species but not *T. ephemeraeformis*. **Bottom center:** Orthologs found in all three species. *Thyridopteryx ephemeraeformis* lacks apyrene sperm, so apyrene classifications are based on the other two species. In both bottom comparisons, the * in the eupyrene partition reflects the fact that a single gene (*Msex2.13702/DPOGS211811/ g11581)* was identified as present in all three species but not found in the sperm proteome of *T. ephemeraeformis*; thus, it is not necessarily conserved in function. For the purposes of counts above, it is considered present in *T. ephemeraeformis*. Image credits: *Danaus plexippus* – Jacobus de Roode, *Manduca sexta* – Andrew J. Mongue, *Thyridopteryx ephemeraeformis* – David Cheng.

Next, we searched for 1-to-1-to-1 orthologs across the full genome annotations for *T. ephemeraeformis, D. plexippus*, and *M. sexta*. We identified 567 strictly sperm orthologs, 146 egg orthologs, and 493 orthologs shared between the two types, based on proteome assignment from *T. ephemeraeformis*. While this approach excludes many protein coding genes, these strictly conserved orthologs represent our highest confidence set of proteins with evolutionarily conserved function (**Figure 2 Middle Right**). We also found significant differences in ortholog conservation between the proteome partitions (X^2^_3_ = 1,564, p < 0.00001), though the biggest difference between groups was genes not identified in either of our proteomes showing far less conservation than those in the sperm or egg proteomes (**Table 1**). Given the larger-than-expected number of annotated genes, it is possible that this result simply represents a higher rate of valid gene models in the proteomes, for which we have additional evidence than in the large bin of genes not found in either the sperm or egg proteomes.

**Table 1.** Ortholog conservation across proteome partitions.

|  | Sperm proteome | Egg proteome | Both | Neither |
| --- | --- | --- | --- | --- |
| 1-to-1-to-1 Ortholog | 567 (11.7%) | 146 (3.0%) | 493 (10.1%) | 3,663 (75.2%) |
| Not conserved | 653 (2.9%) | 203 (0.9%) | 577 (2.6%) | 21,113 (93.6%) |

Separately, we searched for orthology between the *D. plexippus* and *M. sexta* sperm proteomes (**Figure 2 Middle Left**). Limiting our search to 1-to-1 orthologs, we recovered 40 conserved eupyrene sperm proteins, 119 shared between the two morphs, and a single conserved apyrene sperm ortholog (*Msex2.04573/DPOGS215990*). We then intersected these data with the 1-to-1-to-1 dataset and identified 85 proteins found in sperm of all three lepidopteran species. Using the partitions from *M. sexta* and *D. plexippus*, we found 14 eupyrene sperm proteins, 70 shared sperm, and the single apyrene sperm ortholog conserved in the eupyrene sperm of *T. ephemeraeformis*, leaving 24 eupyrene and 44 shared sperm proteins from the other two species missing from *T. ephemeraeformis* (**Figure 2 Bottom**). Note in almost every case missing orthologs were simply not detected between species. One protein (*Msex2.13702*|*DPOGS211811*|*g11581*), however, was strictly conserved in the genomes of all three species, but its expression was not detected in the sperm of *T. ephemeraeformis*. In all other cases, strictly conserved sperm proteins were found to be sperm-specific or shared between sperm and egg proteomes of *T. ephemeraeformis*.

### Characterization of reproductive proteins of interest

For the smaller subset of conserved proteins, we further generated functional annotation via BLAST on NCBI’s protein database. For each protein, we considered the identity of the top hit, any functional annotation, as well as whether or not hits were unique to apoditrysia in Lepidoptera (the group containing *M. sexta* and *D. plexippus*) or more broadly conserved across insects and other animals. We present the full list of these results as supplemental files and here focus on the most interesting patterns and candidates for understanding the evolution of dimorphic sperm.

Starting with 1-to-1-to-1 conserved orthologs, general hits outside of higher Lepidoptera (Ditrysia) were more common than not (**Table 2**). For eupyrene-associated proteins only 7 were specific to Lepidoptera; for those shared between sperm morphs, only 17 were lepidopteran-specific, suggesting that many of these 1-to-1-to-1 conserved orthologs have an even longer history of evolutionary conservation. These included expected functions, including hits to cilia and flagella-associated proteins as well as mitochondrial associated proteins. In terms of patterns of conservation, the most interesting among the conserved orthologs is the single conserved protein in apyrene sperm of the two dimorphic sperm species and the single sperm type of *T. ephemeraeformis*. BLAST searching on NCBI’s protein database recovered hits exclusively to other Lepidoptera, almost all of which are “hypothetical protein[s]” or “unnamed protein product[s]” but the newer annotation of *M. sexta* yielded the functional annotation of “phosphatase 2C and cyclic nucleotide-binding/kinase domain containing protein.”

**Table 2.** Uniqueness of conserved sperm proteins to Lepidoptera.

|  | Unique to Ditrysia | Broader Conservation |
| --- | --- | --- |
| 1-1-1 Conserved | 26 (31%) | 59 (69%) |
| 1-1 Conserved, missing in *T. ephemeraeformis* | 46 (68%) | 22 (32%) |

Turning to sperm proteins lacking in *T. ephemeraeformis*, the proteins in this subset were enriched for lepidopteran-specific and Ditrysian hits, a distinct pattern from that seen in the 1-to-1-to-1 conserved proteins (**Table 2**, X^2^_1_ = 19.36, p = 0.0001). Within this dataset, we identified five trypsin proteinase orthologs. Such proteins are involved in the dissociation of sperm bundles, suggesting potential differences in eupyrene sperm dissociation in *T. ephemeraeformis* compared to the other two species. We also identified three putative seminal fluid proteins unique to the dimorphic sperm proteomes, suggesting a potential role for additional proteomic complexity post-spermatogenesis in these species.

### Sex chromosome identification and composition

Finally, we characterized some basic features of the sex chromosome to understand evolutionary dynamics of reproductive proteins. Karyotyping confirmed that *T. ephemeraeformis* possesses sex chromosome constitution ZZ/Z0 (Figure S1, left). To identify the sex chromosome Z in the assembly, we aligned female reads to the final curated assembly and calculated depth of coverage per scaffold. For all chromosomal scaffolds, the mean coverage depth was 30.9x and the range, with one exception (scaffold_1), was 29.2x – 36.7x. Scaffold_1 had a coverage of 15.8x in the female sample. This matches the expectation that the Z scaffold should have half the coverage of diploid scaffolds in the heterogametic sex and as such we identified it as the Z chromosome. The much smaller debris scaffolds ranged in coverage from 0x to 171x coverage; we did not assign sex linkage to these high-variance, gene-poor sequences. When exploring composition of the sex chromosome, we found a significant difference in the types of protein coding genes compared to the autosomes (X^2^_3_ = 16.2, p = 0.001, **Table 3**), a pattern driven by an increase in sperm-specific proteins and a large decrease in egg-specific proteins on the Z chromosome. Interestingly, we did not see a difference in the overall rate of conservation of orthologs between the Z and autosomes (X^2^_1_ = 0.32, p = 0.571), perhaps because of the very long divergence time between species.

**Table 3.**
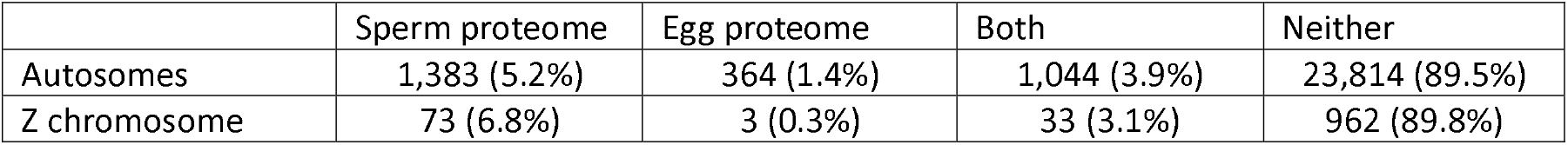
Distribution of reproductive proteins across the *T. ephemeraeformis* genome. Discussion.

We investigated sperm biology of Lepidoptera using *Thyridopteryx ephemeraeformis*, the evergreen bagworm moth, from both cytological and bioinformatics perspectives. In the process, we generated a chromosome-level genome assembly, RNAseq datasets across male and female tissues, a set of gene annotations, and proteomes for sperm and eggs to explore the composition of sperm in this species. Although our focus is on the sperm biology of this species, we first consider new insights for reproductive traits more broadly.

To begin, we found that, like other bagworm moths (Dalíková et al. 2017; Hejníčková et al. 2019), *T. ephemeraeformis* employs ZZ/Z0 sex determination, lacking a female-specific W chromosome found in most Lepidoptera. The Z sex chromosome was enriched for sperm-specific proteins, as expected based on previous study of lepidopteran sperm proteins (Mongue and Walters 2017). Moreover, we also observed a paucity of egg proteins on the Z, a completely novel result to our knowledge. Both of these results align well with predictions from sex chromosome theory (Klein et al. 2021) and empirical validation based on gene expression, which typically finds that the Z is masculinized, *i.e*. having proportionally more male-biased genes, and in some cases, defeminized (having fewer female-biased genes) than the autosomes as well (Mongue et al. 2022; Mongue and Baird 2024; Mora et al. 2024). Whether or not these proteins are evolving more rapidly and/or adaptively on the sex chromosome than the autosomes remains to be tested though, as such patterns are typically (Sackton et al. 2014; Mongue et al. 2022) but not universally (Rousselle et al. 2016; Baird et al. 2025) observed in other insects.

For brevity, and because there is less comparative research on which to draw for lepidopteran eggs, we do not go into detail here about the egg-specific proteins of *T. ephemeraeformis*. We do note that egg proteins showed proportionally less strict orthology between our comparison species than sperm proteins did. We note that *T. ephemeraeformis* diapause and overwinter as eggs (Rhainds et al. 2009), unlike *D. plexippus* and *M. sexta*, which diapause as adults and pupae respectively (Bell et al. 1975; Urquhart 1976). So, in addition to the typical fast evolution of reproductive proteins, there are likely adaptations required to allow eggs to overwinter before development and these newly generated data may serve as a foundation to explore such molecular innovations in the future.

### A new observation of monomorphic sperm and the evolution of apyrene sperm in Lepidoptera

We observed only a single sperm type in *T. ephemeraeformis* and found that morphologically it resembled eupyrene sperm, with obvious nuclear DNA and individual cells bundled together in the seminal vesicle. We found no evidence for apyrene sperm in this species. These results are similar to an earlier report in another bagworm, *Psyche casta*, which found only a very small percentage of sperm were apyrene (~2.6%, Shepherd and Dickinson 2021). This study did not verify identity of apyrene sperm via staining however, so the few reported apyrene sperm may have been misidentified individual eupyrene sperm. Conversely, if apyrene sperm are truly present as a small minority of sperm, it is possible we may have failed to detect them in *T. ephemeraeformis*. Regardless, with estimates ranging from 0 to 2.6%, the sperm biology of bagworm moths stands in stark contrast to better-studied macrolepidoptera, in which apyrene sperm can outnumber eupyrene sperm by 20 to 1 or more (Oberhauser 1989). These observations raise the question of whether bagworms are unique exceptions to broader lepidopteran patterns or more consistent with other early-diverging moths.

Our review of recent observations of apyrene sperm across species suggests the latter. Considering the phylogenetic placement (Regier et al. 2015) of species with reported apyrene and eupyrene sperm counts (Shepherd and Dickinson 2021) suggests that a low ratio of apyrene to eupyrene sperm (<1), verging on no apyrene sperm, is quite common among early-diverging moths. More abundant apyrene sperm appear to evolve in the Ditrysian moths and butterflies after the split from bagworms and other Tineoidea. Further study of early-diverging moths will be required to confirm the consistency of this pattern and more precisely place the increase in apyrene sperm abundance, but for now it raises the question of whether apyrene sperm perform the same function(s) in early-diverging moths and, by extension, if eupyrene sperm have the same limitations there. To begin to answer these questions, we explored patterns of innovation and conservation in sperm proteomes.

### Conservation of shared sperm proteins, variability of morph-specific proteins

We compared the new eupyrene sperm proteome of *T. ephemeraeformis* to the proteomes of apyrene and eupyrene sperm in the obtectomeran Lepidoptera *M. sexta* (Whittington et al. 2015) and *D. plexippus* (Whittington et al. 2017). A previous pairwise comparison found that proteins expressed in both eupyrene and apyrene sperm showed the most proportional conservation between species, followed by those specific to eupyrene sperm, with apyrene-specific proteins showing the least conservation (Whittington et al. 2019). With the *T. ephemeraeformis* sperm proteome, we recovered a similar pattern: the most conservation of orthologs from the set of proteins found in both eupyrene and apyrene sperm of other species, followed by those found only in eupyrene sperm. Then, surprisingly, we found a single protein conserved in the apyrene sperm of *D. plexippus* and *M. sexta* that was also found in the eupyrene sperm of *T. ephemeraeformis*. This protein appears to be unique to Lepidoptera, though its functional significance is uncertain. The only putative functional annotation we recovered was to phosphatase 2C, the function of which is described largely from plants (Rodriguez 1998). Such cross-domain shared amino acid motifs do not imply conserved function, so further work will be needed to study this lepidopteran sperm protein.

Many proteins had more expected hits, however. We recovered 85 strict 1-to-1-to-1 orthologs between the three species’ sperm proteomes, of which 26 had BLAST hits exclusively to Ditrysia. Conversely, 46 of the 68 proteins conserved in *D. plexippus* and *M. sexta* but missing in *T. ephemeraeformis* hit to Ditrysia. This result fits well with a recent broad review of sperm protein evolution, which found that the eupyrene sperm proteomes of *M. sexta* and *D. plexippus* were enriched for lineage-specific proteins compared to other metazoan sperm (Matte et al. 2025). Our results from *T. ephemeraeformis* suggest that early-diverging moth sperm contains more broadly conserved proteins and that lineage-specific innovations are likely unique to (non-Tineoid) Ditrysia, further evidence that the functions and limitations of dimorphic sperm may not be universal in Lepidoptera. This line of thinking is bolstered by considering some of the functional differences recovered between the proteomes. Most intriguingly, we found that 5 of the 68 1-to-1 orthologs conserved in the dimorphic sperm of *M. sexta* and *D. plexippus* but not *T. ephemeraeformis* resemble trypsin enzymes. Trypsin-like enzymes are known to be made by males and transferred to females to break down the extracellular proteins that bundle eupyrene sperm together (Shepherd 1974; Aigaki et al. 1994; Takemura et al. 1996; Friedländer et al. 2001).

### An overlooked feature of dimorphic sperm in Lepidoptera: bundling

While a number insects have sperm superstructures that have been referred to as bundles (reviewed in Dallai et al. 2016), many of these are aggregations of cells that are otherwise unpackaged (e.g. interlocking head and acrosomal pieces with free tails in bushcricket sperm, Viscuso and Vitale 2015). Lepidopteran eupyrene sperm is canonically more intricately packaged: each individual eupyrene sperm cell is enclosed in an inner and outer envelope and groups of eupyrene cells arising from the same cyst in the testis are kept bundled together by fibrous extra-cellular proteins (Friedländer et al. 2001). These extracellular proteins appear to derive from sperm structures termed lacinate appendages (André 1959) that present as extensions of the cytoplasm of mature sperm in the testes, but change conformation when sperm leave the testes to form the extra-cellular proteins that bind together eupyrene sperm bundles in the seminal vesicle (Phillips 1970). *In vitro* treatment of sperm bundles with trypsin digests the extracellular proteins until they have dissociated into individual eupyrene cells that only then become motile (Friedländer et al. 2001). In contrast, apyrene sperm lack lacinate appendages in the testes (Phillips 1971), are not bundled in the seminal vesicle, and become motile without the need for a trypsin treatment (Friedländer et al. 2001).

As shown here in *T. ephemeraeformis*, eupyrene sperm are still bundled in the male reproductive tract, but, anecdotally, the bundles dissociated more readily during mounting and staining than expected based on other lepidopteran sperm. Because adult bagworms of both sexes are short-lived (with lifespans of roughly 48 hours, Kaufmann 1968)and thus do not store sperm for as long after spermatogenesis or mating (Rhainds et al. 2009), it may be that their eupyrene sperm do not require as robust of bundling as other longer-lived species and thus do not require as much enzymatic assistance to dissociate and become motile.

### How did the essential function(s) of apyrene sperm evolve?

As discussed above, apyrene sperm likely perform multiple roles and like most other traits are likely to have adaptive lineage-specific roles, such as helping avoid sperm competition in polyandrous species (Wedell and Cook 1999; Swallow and Wilkinson 2002). Here, however, we wish to focus on their fundamental role as essential components of successful fertilization. In the simplest possible terms, the function of fertilizing sperm is to deliver DNA to the egg. In other words, it is a cell type that must reach a target in an extracellular medium. For example, some fish sperm actively responds to chemical cues in female ovarian fluid to move towards the eggs in a species-specific manner that leaves heterospecific sperm moving in circles (Butts et al. 2012). When asking how apyrene sperm are required for fertilization, it is necessary to understand the movement dynamics of sperm in the female reproductive tract in Lepidoptera.

There are relatively few studies on this topic, but those that do exist tend to be very detailed. Observation of recently mated female *M. sexta* suggests roles for both active motility and passive transport; namely, both sperm types exit the male spermatophore under their own power, but are apparently transferred to the spermatheca by muscular contractions of the female seminal duct (Hague et al. 2021). Hague et al. report that apyrene sperm are already motile within the spermatophore at the end of copulation and that eupyrene sperm only slowly acquire motility as bundles dissociate into individual cells over a period of several hours. Consequently, apyrene sperm begin to reach the spermatheca long before eupyrene sperm do. More specifically, the spermatheca of *M. sexta* is a two-lobed organ, with a shorted bulb called the lagena and a longer portion known as the utriculus and Hague et al. describe apyrene sperm initially collecting in the utriculus but then arriving in the lagena along with eupyrene sperm. In the longer term, eupyrene sperm are stored in the utriculus but apyrene sperm begin to disappear from both parts of the spermatheca (Hague et al. 2021).

By knocking out apyrene sperm production in *B. mori*, Sakai et al. showed that the remaining eupyrene sperm never reach the spermatheca in matings with these mutant males, despite becoming motile in the bursa copulatrix (2019). Much earlier work using triploid male *B. mori*, which do not produce functional eupyrene sperm, showed that even in their absence, apyrene sperm still reach the spermatheca (Sugai and Sugita 1976). Putting these observations together, it seems likely that apyrene sperm assist in the movement of eupyrene sperm throughout the female reproductive tract. This assistance could be as direct as physically moving eupyrene sperm, for example through the semiporous contents of the spermatophore, or by maintaining female physiological response of peristalsis long enough for eupyrene sperm to reach the base of the seminal duct and be transferred under female control. More work will be required to refine or refute these hypotheses, but it is worth stating clearly, as there has been little synthesis between the distinct works discussed above.

Next, we consider how evidence from *T. ephemeraeformis* fits with the observations above. In step with other morphological idiosyncrasies, *T. ephemeraeformis* have very atypical lepidopteran reproductive traits, the three most relevant differences being (1) the spermatheca of females is single chambered, with that single chamber morphologically more resembling the bulbous lagena, (2) male bagworms do not transfer a spermatophore to females (Williams 1941; Kumpulainen 2004), and (3) the female seminal duct is highly elongated (Davis 1964). The first of these is consistent with an absence or paucity of apyrene sperm as the sperm storage organ no longer has two compartments for storage of different sperm types. The second observation is as well; with males transferring only sperm and seminal fluid to females, presumably because they do not live long enough to digest any nutrient nuptial gift, eupyrene sperm do not have to navigate exiting the spermatophore before entering the female reproductive tract. The third observation would seem at odds with at least part of hypothesis above though. If sperm require female muscle contractions to traverse the seminal duct and apyrene sperm initiate or maintain that response, how would *T. ephemeraeformis* eupyrene sperm make their way through an exceptionally long seminal duct without the support from apyrene sperm?

Two possibilities present themselves. First, the function of the seminal duct could be independent of the presence or absence of apyrene sperm, perhaps assisting in movement in the spermatophore is the key function of Ditrysian apyrene sperm. Indeed, both *T. ephemeraeformis* and *P. casta* have few to no apyrene sperm, but *P. casta* has a drastically shorter seminal duct (Davis 1964), demonstrating that these two traits are not strongly correlated. It is interesting to note that as barely mobile adults, females seemingly have little control over mate choice. In species with highly reduced females like *T. ephemeraeformis*, females put out pheromone calls to males, which then insert their telescoping abdomens into the bag to mate (Rhainds et al. 2009). In other words, with little opportunity for precopulatory choice, post-copulatory cryptic choice may be more relevant for *T. ephemeraeformis* (Eberhard 1996). In contrast to the overall trend for monandry in bagworms (Rhainds et al. 2009), Jones describes occasional double matings in *T. ephemeraeformis* (1927). A long sperm-transporting organ certainly would give the opportunity, though on what basis females might choose remains to be seen.

Inbreeding avoidance has been proposed as basis for cryptic choice in another Lepidopteron, but subsequent testing found no evidence for post-copulatory inbreeding avoidance (Mongue et al. 2015), despite a clear cost in the form of inbreeding depression (Mongue et al. 2016). Moreover, male and female larvae that hatch at the same time develop at different rates, such that males mature and disperse before females mature (*i.e*., protandry, Rhainds 2013). This premating dynamic may be sufficient to minimize inbreeding, but other factors may be relevant to cryptic choice.

The second possibility is that the lack of apyrene sperm may not imply the lack of the functional role canonically fulfilled by apyrene sperm. As tenuous evidence to support this possibility, we note that one apyrene-specific protein was maintained in a strict 1-to-1 orthologous relationship between *D. plexippus* and *M. sexta* and that protein is also found in the single, eupyrene sperm type of *T. ephemeraeformis*. This protein has little functional annotation, so it is pointless to speculate as to its molecular role. Still, it is at least unique to Lepidoptera and makes an appealing target for functional genomic knockout/knockdown experiments in the future. Alternatively, the core functionality of apyrene sperm could be found in some of the 1-to-many or many-to-many orthologs, or even in proteins that evolve too quickly to detect conservation between these diverged lineages. If apyrene sperm are interacting with female physiology, they may be rapidly evolving in concert with female reproductive tract proteins, as seen in some marine invertebrates (Swanson and Vacquier 2002; Zigler et al. 2005). *Thyridopteryx ephemeraeformis* last shared a common ancestor with *D. plexippus* and *M. sexta* roughly 150 million years ago, during the Jurassic, and the latter two diverged roughly 100 million years ago (Kawahara et al. 2019), so there has been ample time for such change between lineages.

### Why were apyrene sperm gained in Lepidoptera?

Regardless of current instrumentality to fertilization, originally apyrene sperm had to have evolved in an ancestor with functional monomorphic eupyrene sperm (*sensu lato*, nucleated sperm cells). It is worth considering the scenarios under which such evolution could be favored. Eupyrene sperm production typically ceases late in male larval or early pupal development and apyrene sperm production begins in pupal males, often continuing into adult moths and butterflies (Friedlander 1997; Friedländer et al. 2005). In this way, Lepidoptera resemble other insects for which (fertilizing) sperm production ceases in adulthood. This developmental life history is often associated with short-lived, often non-feeding adults (Heming 2018). And indeed, in at least one species of caddisfly (Trichoptera), the sister order to Lepidoptera, spermatogenesis appears to cease prior to adult eclosion, as the testes contain only mature sperm at that point (Costa et al. 2024, but note that sperm morphology is odd in caddisflies, with largely immobile sperm, so spermatogenesis may be idiosyncratic). Based on these observations, apyrene sperm production seems to be a novel extension of spermatogenesis into the adult insect which may be associated with longer life. Some have proposed that the evolution of flowering plants and subsequent shift to adult Lepidoptera feeding on nectar created new opportunities for adult niches and longevity (Stekolnikov and Korzeev 2007) and more recent evolutionary genetic work has shown Lepidoptera acquired many new gene families that may have helped them implement new strategies across life history traits from egg-laying to detoxification around this time (Weng et al. 2025).

In a monomorphic sperm ancestor, it is difficult to imagine apyrene sperm immediately evolving as a necessary capacitator of fertilization. Instead, apyrene sperm production may have started in a supporting role, such as a way to avoid sperm competition as adult Lepidoptera began to live longer and the risk of female remating increased. Previous work has failed to recover a signal for stronger positive selection on apyrene sperm proteins in more polyandrous species (Mongue et al. 2019), but observational work suggests that females do delay remating longer when they receive more apyrene sperm (Oberhauser 1989; Wedell and Cook 1999). Thus, while apyrene sperm may not be sophisticated agents of sperm competition, their presence may help avoid such competition before it can occur. Adult feeding and longer lifespans would provide males with the resource to produce more apyrene sperm. The relationship between adult lifespan and remating are not so simple, however.

Non-feeding, short-lived Saturniid moths can still be polyandrous (Morton 2009), and the longer-lived Sphingid *M. sexta* is generally monandrous (Snow et al. 1974). Moreover, both of these groups produce large quantities of apyrene sperm (Friedlander 1997; Shepherd and Dickinson 2021). Considering these observations, short lifespan itself is not a good predictor of polyandry or the presence of apyrene sperm. The relative absence in bagworms then cannot be explained by short lifespans alone. Where mating rates have been examined, female bagworms show a high rate of mortality before mating, and when successful are generally observed to mate only once (as reviewed in Rhainds et al. 2009). Because females of most species are barely motile and do not leave their larval cases, the risk of sperm competition may be generally low across most of the family, excepting a confirmed low rate of polyandry in *Thyridopteryx* (Jones 1927). These patterns are broadly consistent with the observation that apyrene sperm are low abundance in the short-lived early diverging moths and further suggests their role in those taxa may be distinct.

Furthermore, with only functions in sperm competition, however, early apyrene sperm would not be strictly necessary for successful fertilization. But as adult lifespan increased, so too would the time between eupyrene spermatogenesis and copulation. As shown both here in *T. ephemeraeformis* and in other Lepidoptera (Whittington et al. 2019), mature eupyrene sperm are stored in quiescent bundles, held together by extra-cellular proteins. It may be that longer adult lifespans require more robust bundles to efficiently store and preserve eupyrene sperm until mating. The fact that the long-lived *M. sexta* and *D. plexippus* sperm proteomes both include protein digesting enzymes that are missing from the sperm of the short-lived *T. ephemeraeformis* suggests as much. To be sure, this hypothesis is highly speculative, and it is impossible to verify the sperm biology of ancestral Lepidoptera, but hopefully future study of early-diverging moths can offer additional evidence for or against this scenario.

## Conclusions

We have described the sperm biology of the evergreen bagworm, *T. ephemeraeformis*, an informative outgroup to better studied Lepidoptera. In the process, we have reviewed evidence for the roles of apyrene sperm and identified candidates for further study. More specifically, we have generated a number of bioinformatic resources that will be useful in studying all aspects of the unique biology of bagworms. Putting previous observations and new data together, we show evergreen bagworms possess only eupyrene fertilizing sperm. Moreover, the eupyrene of *T. ephemeraeformis* appears somewhat distinct to that of later-diverging, longer-lived moths at the molecular level. In particular bagworm sperm appears to contain more deeply conserved sperm proteins and fewer lepidopteran-specific proteins, including a lack of trypsin-like enzymes used to digest the proteins that bind and immobilize individual eupyrene sperm cells into bundles. Given that these bundles must break down for successful fertilization, we infer that they are not as strongly constructed, because they do not require enzymatic assistance for degradation.

Finally, *T. ephemeraeformis* has few to no apyrene sperm, a condition that appears common in other early-diverging moths. We conclude that apyrene sperm was not immediately necessary for fertilization when it first evolved and likely still is not in these taxa. Based on observed patterns of sperm movement in the female reproductive tract under both natural and experimentally manipulated conditions, apyrene sperm seem likely to govern the movement of eupyrene sperm out of the spermatophore and/or bursa copulatrix. Bagworm moths lack spermatophores and may have less tightly bound eupyrene sperm, and as such do not need apyrene sperm to assist in the movement of eupyrene sperm.

## Acknowledgments

We thank the members of the Mongue lab for reading and offering feedback on drafts of this manuscript. Thanks to Dr. Jacobus de Roode and David Cheng for use of insect images. Special thanks to Dr. James Walters for help in collecting bagworms remotely and continued support of a project that began seven years ago. Thanks to Dr. Hiroki Sakai for useful conversations around dimorphic sperm in *Bombyx mori*. This project was funded by NSF DEB 2426250 and a previous NSF DEB 1701931 Doctoral Dissertation Improvement Grant award to AJM.

## Notes

### Competing Interest Statement

The authors have declared no competing interest.

