## Supplemental figures and tables for "Insights from the lack of an enigmatic trait: monomorphic sperm in evergreen bagworm moths and the evolution of sperm dimorphism"

**Supplementary methods and results**

*Karyotyping*

Chromosome preparations for cytogenetic analyses were made by spreading technique from gonads of fourth to fifth instar larvae of both sexes following Šíchová et al. (2013). Female and male genomic DNA were extracted from larvae using the HMW MagAttract Kit (Qiagen, Hilden, Germany) and amplified using the illustra GenomiPhi HY DNA Amplification Kit (GE Healthcare, Milwaukee, WI), both following the manufacturer's protocol. To generate a probe for genomic *in situ* hybridization (GISH), female DNA was labelled by nick translation with Cy3-dUTP (cyanine 3-deoxyuridine triphosphate; Jena Bioscience, Jena, Germany) following Kato et al. (2006) with a 3.5-hour incubation at 15°C. Male competitor DNA was fragmented by incubation 20 min at 99°C. GISH was performed following the protocol of Yoshido et al. (2005). For each slide, the hybridization cocktail contained 300 ng of female labelled probe, 3 μg of male competitor DNA, and 25 μg of salmon sperm DNA.

The same chromosomal preparations were also used to localize arrays of major ribosomal RNA genes (rDNA). For fluorescence *in situ* hybridization with the rDNA probe, a fragment of the *Cydia pomonella* 18S rDNA was amplified following Fuková et al. (2005) and labelled by nick translation with biotin 16-dUTP (Roche, Basel, Switzerland) following Kato et al. (2006), with a 1-hour incubation at 15°C. Chromosomal preparations were re-probed as described in Nguyen et al. (2013). For each slide, the hybridization cocktail contained 30 ng of biotin-labelled 18S rDNA probe and 25 μg of salmon sperm DNA. The signal was detected using Cy3-conjugated streptavidin (Jackson ImmunoResearch Laboratories Inc, West Grove, PA, US) and enhanced with biotinylated anti-streptavidin (Vector Labs. Inc, Burlingame, CA, USA). The enhanced signal was amplified again with Cy3-conjugated streptavidin, following Fuková et al. (2005).

Preparations were counterstained with 0.5 mg/mL DAPI (4,6-diamidino-2-phenylindole; Sigma-Aldrich, St. Louis, Missouri, USA) in DABCO antifade (1,4-diazabicyclo[2.2.2]octane; Sigma-Aldrich, St. Louis, Missouri, USA). Results were observed in the Zeiss Axioplan 2 Microscope (Carl Zeiss, Oberkochen, Germany) and documented with an Olympus CCD Monochrome Camera XM10 with the cellSens 1.9 digital imaging software (Olympus Europa Holding, Hamburg, Germany). Images were pseudo-colored and superimposed in Adobe Photoshop CS3.

*Cytogenetics of* Thyridopteryx ephemeraeformis

We performed karyotyping and *in situ* hybridizations using genomic and rDNA probes to learn about the genome structure of *T. ephemeraeformis* from a sequence-independent approach. In mitotic nuclei prepared from gonads of *T. ephemeraeformis* larvae, we counted 62 chromosomes in the male preparation and 61 chromosomes in the female (Figure S1a,c), which is in agreement with a previously reported absence of W chromosome in representatives of the family Psychidae (Hejnickova et al. 2019). Genomic *in situ* hybridization, which is standardly used for detection of the W sex chromosome (e.g. Yoshido et al. 2005), produced strong and weak signals on four and two chromosomes in the female mitotic complement, respectively. However, GISH produced the same hybridization pattern also in male mitotic complements (Figure S1c), which further supports absence of the W chromosome. Additional GISH signals could correspond to arrays of repeats (Picq et al., 2018). Hence, we tried to test co-localization between GISH signals and arrays of genes for major ribosomal RNAs (rDNA). Fluorescence *in situ* hybridization with a probe derived from 18S rDNA highlighted both ends of two mitotic chromosomes in both male and female complements (Figure S1a,c). However, these were different from chromosomes bearing the GISH signals (Figure S1a,c).

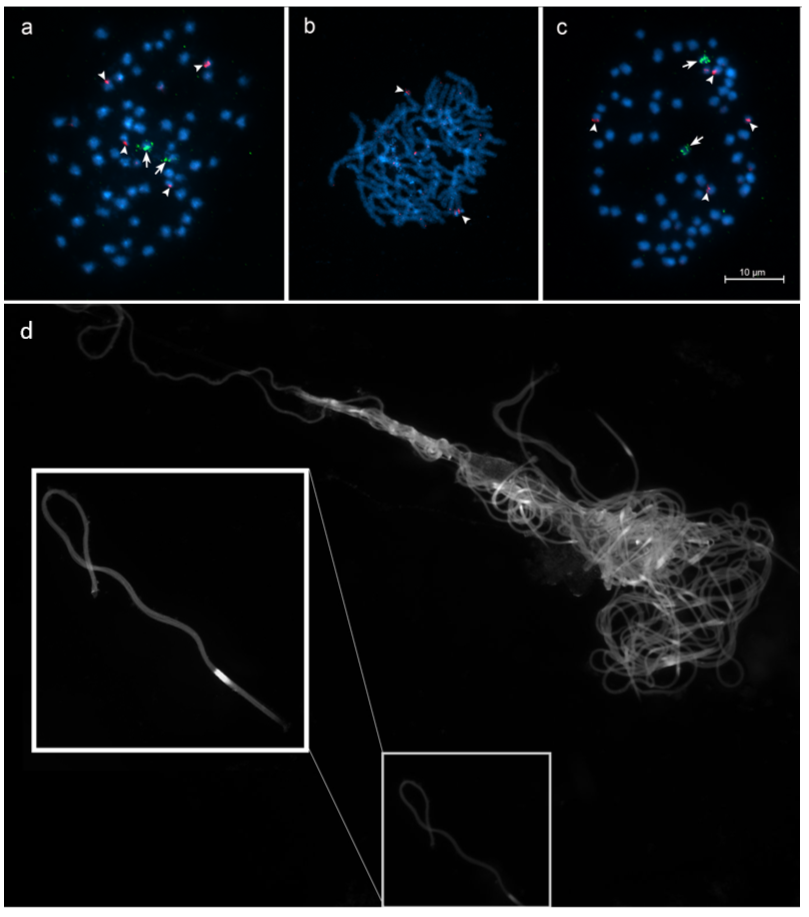

**Figure S1.** Cytogenetic characterization of *T. ephemeraeformis*. **Top:** Chromosomes were stained by DAPI (blue). Arrows mark the rDNA signal (green). Arrowheads mark the female genomic signals (red). The diploid chromosome count is 61 for females (**a**) and 62 for males (**b**), suggesting that the system is ZZ/Z0 with no W chromosome. **c:** Further evidence from Genome *in situ* Hybridization (GISH) in female pachytene nuclei revealed no large portions of the genome unique to females (red), further confirming a lack of W chromosome. Scale bars represent 10μm. **Bottom:** Initial (2018) observation of an apparent lack of apyrene sperm. Whole seminal vesical content of an adult male under fluorescence (**d**). Cells are stained with DAPI which appears as a brighter spot where it contacts nuclear DNA. All sperm cells appear bundled with nuclear DNA regions in the head. A magnified view of a single eupyrene bundle is shown in the inset (**box**).

| Sample | NCBI Accession or repo link |
| --- | --- |
| DNA | |
| PacBio HiFi reads | SRR32029783 |
| Hi-C reads | SRR32029781 |
| Female short-read sequencing | SRR32029782 |
| Genome assembly | JBLDYN000000000 |
| RNA | |
| Adult male head | SRR32037186, SRR32037185 |
| Adult female head | SRR32037189 |
| Adult male thorax | SRR32037196, SRR32037182 |
| Adult male testes | SRR32037191, SRR32037190 |
| Adult male accessory gland | SRR32037197, SRR32037198 |
| Adult female ovaries | SRR32037194, SRR32037193 |
| Adult female ovipositor | SRR32037195 |
| Male pupa | SRR32037184, SRR32037183 |
| Female pupa | SRR32037187, SRR32037188 |
| Unsexed larval silk gland | SRR32037192 |
| Annotation | |
| Gene annotations | <https://github.com/amongue/ThyridopteryxGenome/> |
| Protein | |
| Sperm proteome hits to annotations | <https://github.com/amongue/ThyridopteryxGenome/> |

**Table S1. Archiving location for data generated in this study.** Accessions beginning with SRR can be found on NCBI. Additional resources including annotation fastas and analysis scripts can be found on the git repo for this project.

| Assembly Statistics | Primary HiFi Assembly (Hifiasm) | Hi-C Scaffolded Assembly (YaHS) | Final Curated Assembly  (Juicebox) |
| --- | --- | --- | --- |
| Total length | 818,398,161 bp | 818,402,761 bp | 818,402,861 bp |
| Sequence count | 154 | 120 | 123 |
| N50 | 19,951,357 bp | 28,623,661 bp | 28,623,661 bp |
| L50 | 16 | 12 | 12 |
| GC content | 34.18% | 34.18% | 34.18% |
| Percent assigned to chromosomal scaffold | NA | NA | 97.9% |
| BUSCO completeness  (lepidoptera_odb10, n=5,286) | C:95.7%[S:93.8%,D:1.9%],F:0.5%,M:3.% | NA | C:95.7%[S:93.9%,D:1.8%],F:0.5%,M:3.8% |

**Table S2. Statistics for genome assembly at each stage of the assembly process.**
